# Size spectra in freshwater streams are consistent across temperature and resource supply

**DOI:** 10.1101/2024.01.09.574822

**Authors:** Vojsava Gjoni, Justin P. F. Pomeranz, James R. Junker, Jeff S. Wesner

## Abstract

The study explores the individual size distribution (ISD) pattern in ecological communities, characterized by a negative correlation between individual body size and abundance (N ∼ M^λ^). The parameter λ denotes the rate of decline in relative abundance from small to large individuals. Despite known influences of temperature and resource availability on body size, their effects on λ remain diverse. Leveraging data from 2.4 million individual body sizes in continental freshwater streams, the research the hypothesis that λ varies as a function of temperature and resource supply. Surprisingly, despite varied environmental conditions and complete species turnover, minimal variation in λ (mean = −1.2, sd = 0.04) was observed, with no discernible impact from temperature or resource supply. The unexpected λ value of −1.2 suggests a higher-than-expected relative abundance of large individuals, challenging assumptions of metabolic scaling at 0.75 and implying large subsidy inputs to large predators. Simulation and mesocosm experiments support a metabolic scaling coefficient of ∼0.4 for freshwater macroinvertebrates. The findings underscore remarkable consistency of individual size distributions in freshwater streams, likely driven by shallow metabolic scaling and large subsidies to large consumers.

---

A common pattern in ecological communities is the negative relationship between individual body size and abundance, described by a power law N ∼ M^λ^ ^1–4^. This pattern is known as the individual size distribution (ISD)^5^, where λ is the rate of decline in relative abundance from small to large individuals. The ISD is a powerful tool for assessing changes in spatiotemporally and taxonomically disparate ecosystems ^3,6–8^. This is because λ is thought to vary as a function of trophic transfer efficiency, predator-prey mass ratio, and metabolism-mass scaling, universal processes across ecosystems ^4,9–11^. Temperature and resource availability have been shown to alter λ, but the direction and magnitude of their effects are varied. Here, we leverage 2.4 million individual body sizes from freshwater streams spanning natural gradients of temperature and resource availability to estimate changes in λ at a continental scale. Despite broad environmental conditions and complete species turnover, we find little variation in λ (mean = −1.2, sd = 0.04) and no change with temperature or resource supply. The value of λ = −1.2 represents higher than expected relative abundance of large individuals. The only way to achieve this is by relaxing the assumption that metabolic scaling is 0.75 and by assuming subsidy inputs to large predators. We support these hypotheses with a simulation study and mesocosm experiment that suggests a metabolic scaling coefficient of ∼0.4 for freshwater macroinvertebrates. Our results emphasize remarkable uniformity in the prevalence of larger individuals within freshwater streams, persisting across both spatial and temporal dimensions. A critical next step is to understand the mechanisms for upholding the constancy of λ under varying environmental conditions.

## Main

Earth supports ∼550 gigatons of living carbon biomass ^12,13^ and a fundamental challenge for ecologists is to understand how that biomass is distributed across scales from individuals to ecosystems ^14^. The individual size distribution (ISD) is a common method for describing how biomass is distributed across individuals within communities ^5,15^ and is driven by the flow of energy through food webs ^3,16–18^. Size-based assessments, such as the ISD, are emerging as a powerful tool to bridge individual-level physiological processes to ecosystem-level patterns ^18^, complementing more traditional taxonomic or trophic approaches ^3,19^. This is because many fundamental aspects of an organism’s biology are controlled by body size, including metabolic rate, life history, diet breadth, and trophic position ^5,20,21^. Therefore, changes in the ISD reflect changes in fundamental attributes of a community, providing a measure of variation in ecosystem structure and function (Fig. 1^1–4^). The ISD is described by a power law, *N ∼ M ^λ^*, where *N* is abundance and *M* is individual body mass. Metabolic scaling theory predicts that the exponent *λ* is generated by interactions of three ecosystem-level variables: trophic transfer efficiency (α), predator-prey mass ratio (β), and metabolic scaling (γ)^4,9–11^, such that:

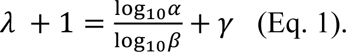

**Fig. 1.**
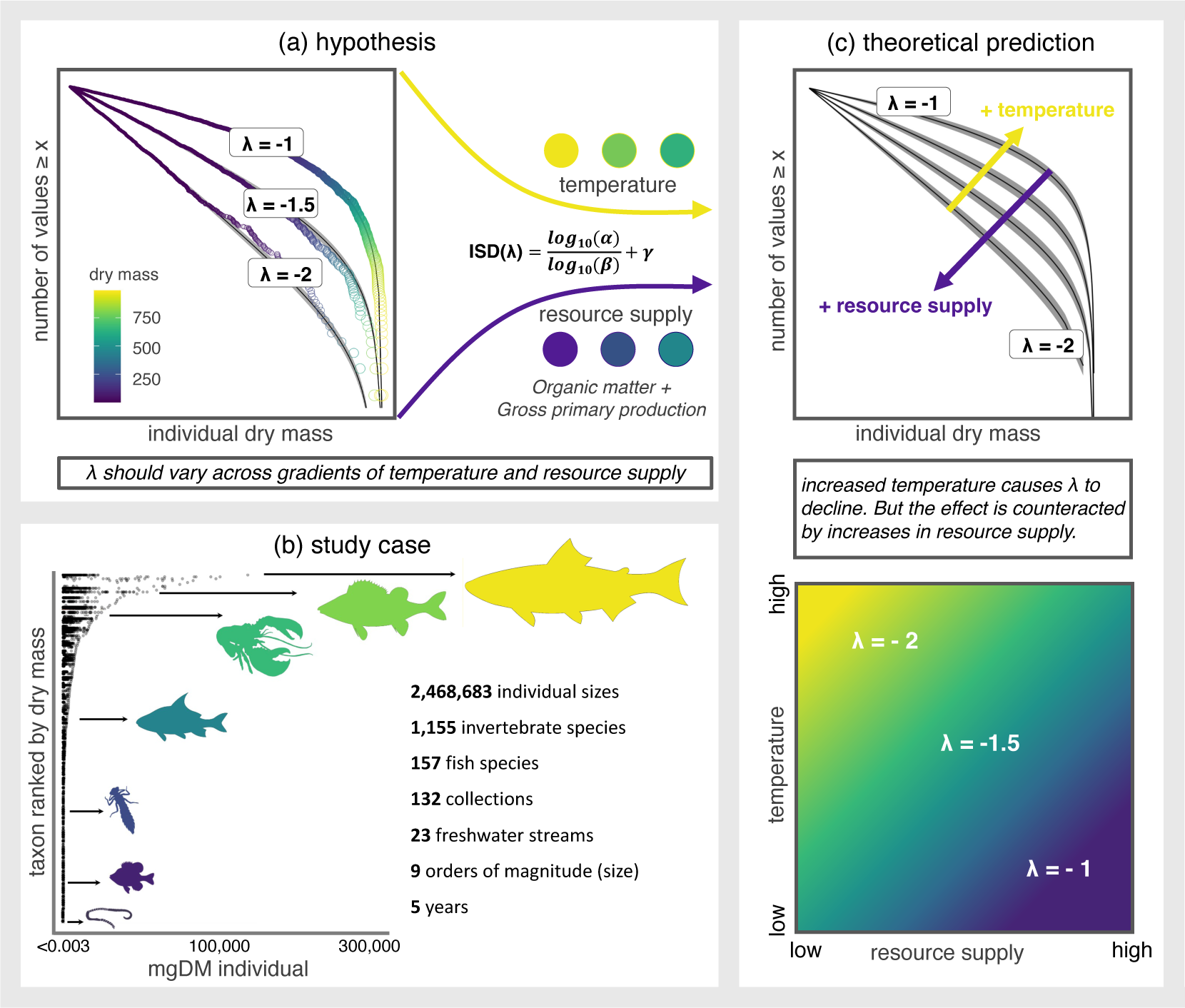
The theoretical basis for Individual Size Distribution (ISD) scaling in ecosystems is underpinned by several empirical studies and theories in ecology. a. At the community level, λ consistently exhibits a negative pattern between −1 and −2. While the individual impacts of temperature and resource supply on the ISD have been extensively investigated, their combined or synergistic effects remain largely unexplored. In this study, we propose that λ varies in response to temperature and resource supply. Our hypothesis is that temperature and resource supply interact to shape λ across food webs. b. Our study centers on the NEON continental-scale data collections (e.g., 2,468,683 individual sizes), from both fish (157 species) and macroinvertebrates (1,155 species). c. Theory predicts that λ should decline with increasing temperature. However, these effects can be mitigated by an increase in resource supply, as demonstrated by previous studies and theory^4,20,30^. The range of λ spans from ∼ −1 to −2, where smaller λ values (e.g., −2, represented by dark purple, indicating high resource supply) signify less efficient trophic transfer, supporting relatively fewer large organisms compared to ecosystems with higher λ values (e.g., −1.5, −1, represented by green to yellow colors, indicating high temperature).

Given typical values of α, β, and γ, the exponent λ tends to be negative, ranging from about −1 to −2, underscoring a remarkably consistent ecological pattern across Earth’s diverse ecosystems ^3,6–8^. Owing to the apparent consistency of the ISD, body size distributions have been suggested as a “universal indicator” of ecological status^18^. Consequently, ecologists have increasingly used it to indicate fundamental changes in community structure and ecosystem function in response to anthropogenic impacts, including over-fishing and environmental pollution^8,16,22^.

Experimental and empirical results have also shown that λ varies in response to temperature, though the magnitude and direction of change is inconsistent. Increasing temperature is predicted to favor smaller organisms^23,24^ leading to more negative (or “steeper”) values of λ and this has been observed empirically^25–28^. However, λ has also increased (“shallower”) in response to temperature^4^ or had no response^29^. One explanation for the inconsistent response of λ to temperature is compensatory effects of resource supply at the base of the food web^4^. Increasing temperature induces higher metabolic costs, leading to reduced trophic transfer efficiency and a subsequent selection for smaller body sizes. But these effects can be counteracted by increases in resource supply^4,20,30^. However, how temperature and resources interact to affect λ is almost completely overlooked^31^.

Testing how λ responds to temperature and resource supply at the macroecological scale is logistically challenging because it requires data-intensive measures of individual body sizes, accurate (daily or sub daily) temperature measures, as well as estimates of resource supply (gross primary production, allochthonous subsidies)^4,32^. Here, we overcome these challenges by using data from the National Ecological Observatory Network (NEON). NEON is a continental scale ecological sampling program that uses standardized, automated sensor measurements coupled with observational field data and biological collections repeated across seasons (spring, summer, autumn), years (from 2015 to present), and space (across North America). Specifically, we examined the response of 133 ISD’s using individual fish and macroinvertebrate body sizes collected from 22 freshwater streams in North America that varied by 25°C in mean annual temperature and by orders of magnitude in resource supply (i.e., gross primary production and litter input).

**Fig. 2.**
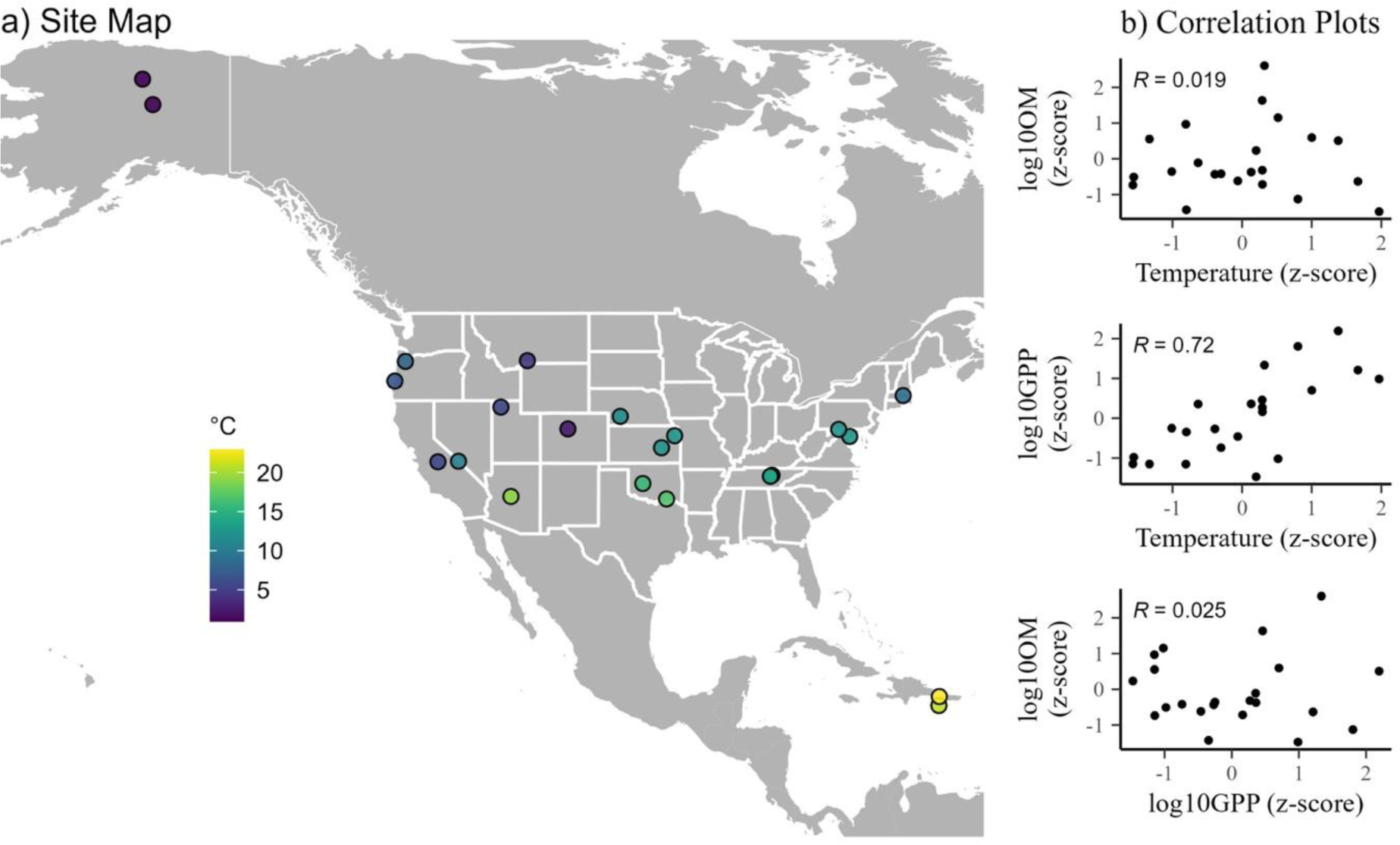
Variation in temperature and correlations of temperature, gross primary production (GPP), and organic matter (OM) across NEON stream sites (n = 22). a. NEON sampling sites span a wide range of mean annual temperatures (1 to 24°C). Each sampling site is a point on the map, with cooler temperatures indicated by shades of blue and warmer temperatures depicted by shades of yellow. b. Variation in environmental variables. Values for temperature and GPP are annual means. Values for organic matter represent standing stocks averaged across 2-4 samples per site.

## Consistent λ’s across temperature and resource supply gradient

Across the 133 samples, *λ* ranged from −1.47 to −1.14 (posterior median) with an average of −1.22 (Fig. 3a; 95% CrI: −1.24 to −1.2). There was no evidence that *λ* varied with temperature (Fig. 3a; marginal slope: −0.002 (95% CrI: −0.03 to −0.02), posterior median (95% CrI), gross primary production (Fig. S1; marginal slope: −0.0002 (95% CrI: −0.03 to 0.03)), organic matter (Fig. S1g; marginal slope: 0.004 (95% CrI: −0.01 to 0.03)), or their interactions (Fig. 3b). These results diverge from predictions of metabolic scaling theory, which predicts λ to decline at elevated temperatures^26^. The theoretical prediction arises from the tendency of body size to decline at higher temperatures, as individual energy demand increases, unless there is a compensatory increase in resource supply^26,31^. If such a compensatory increase explained these results, one would anticipate a negative temperature effect in multiple regression after adjustment for resource supply. No such effect was found (Fig. S1). In fact, environmental variables were not reliably associated with λ under any model formulation, including univariate, two-way, and three-way interactions (Fig. S1). Despite extensive testing, our findings indicated that λ only marginally decreases with rising temperature, specifically under circumstances characterized by high gross primary productivity (GPP) and high organic matter (OM). It is noteworthy, however, that this combination of temperature and resource conditions are extremely rare in NEON streams (Fig 3c).

**Fig. 3.**
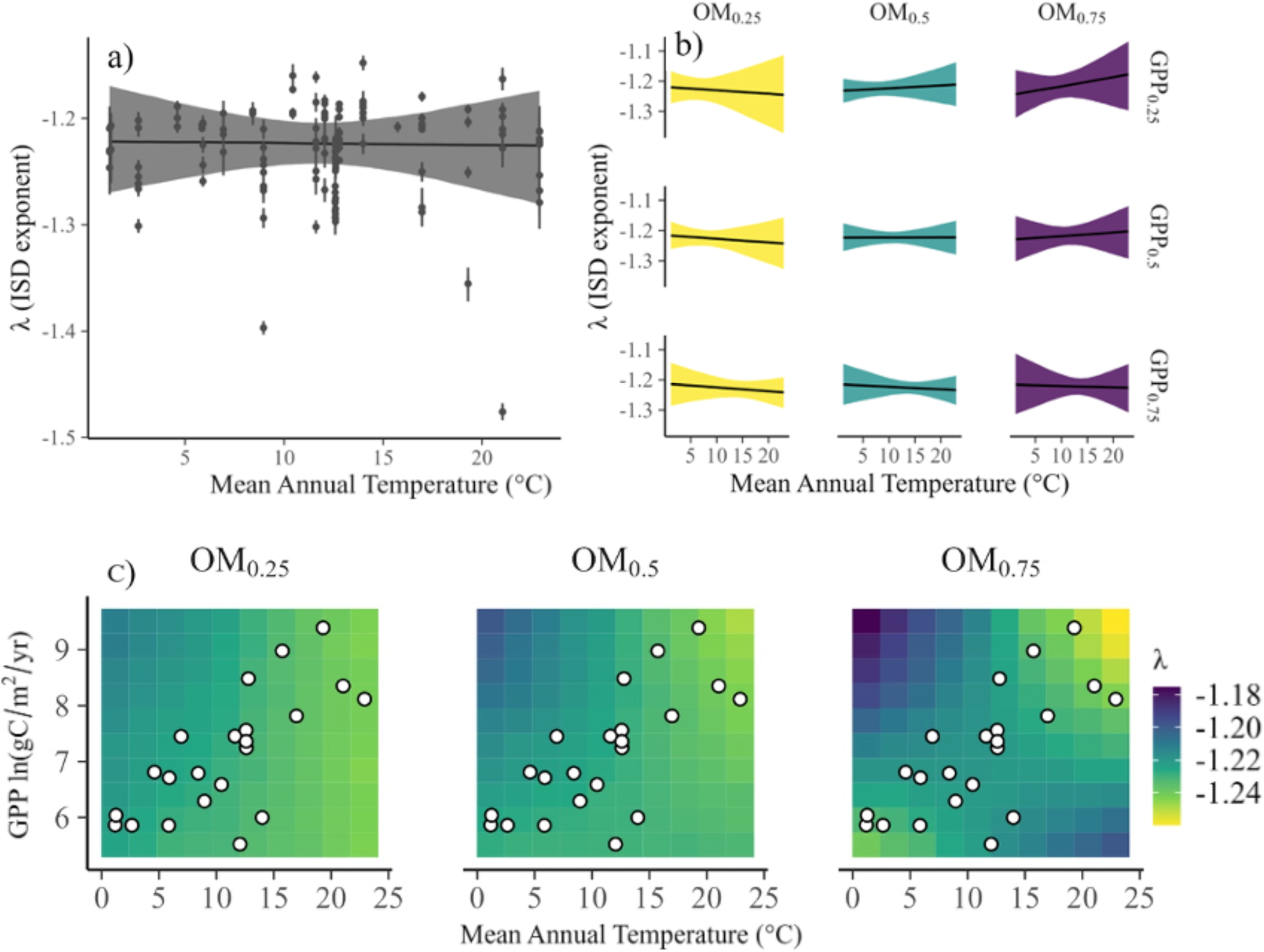
Relationships of the ISD across mean annual stream temperature, annual gross primary production (GPP), and organic matter (OM). a. Mean annual temperature is not related to the ISD exponent (marginal relationship from a multiple regression model). b. Temperature causes no change in ISD across 25^th^, 50^th^, and 75^th^ quantiles of organic matter (OM; facet columns) and gross primary production (GPP; facet rows). The temperature range (23°C) covers ∼90% of the global temperature gradient in stream ecosystems (∼25°C)^41^. In addition, OM estimates of our data are consistent with the two order of magnitude range in streams of North America^42^, as well GPP estimates, with a range from 300 to 11000 g C m^−2^ yr^−1^, that is more than twice the observed GPP in North America^43^. c. Using our model predictions, λ is predicted to decrease slightly with increasing temperature only under conditions of high GPP and high OM. But, as shown by the white dots representing empirical measures, these combinations of conditions do not exist in our data set. Conversely, decreasing temperature leads to an increase in λ, but again, exclusively under high GPP and high OM conditions. No changes in λ were observed under other environmental conditions; the value of λ remained constant.

## λ is larger than expected from theory

While it is clear that λ was invariant to environmental conditions, it is less clear why NEON streams converged to a λ of −1.22 as compared to the theoretically predicted value of −1.95. For example, we can use Eq. 1 to predict λ using standard values of trophic transfer efficiency (α), predator prey mass ratio (β), and metabolic scaling (γ). In practice, γ is assumed to be 0.75, α = 0.1, and β = 10^4^ ^11,20^. Using those values and solving Eq. 1 generates a prediction of *λ* = −1.95, far lower than our empirical estimate of −1.2. The metabolic scaling coefficient γ is particularly important as it is commonly assumed to be fixed at −0.75^11,20^. Under this assumption, *λ* = −1.2 can only be achieved by adjusting trophic transfer efficiency (α) or the predator-prey mass ratio (β). To examine possible values of these parameters in our samples, we used Monte Carlo simulation in which α and β were randomly drawn from probability distributions based on published values from the literature (Table S1). As shown in Fig. 4a, the most extreme combinations *α* and *β* yield *λ*’s no higher than about −1.6, slightly smaller than the smallest empirical value in our samples (−1.47).

**Fig. 4.**
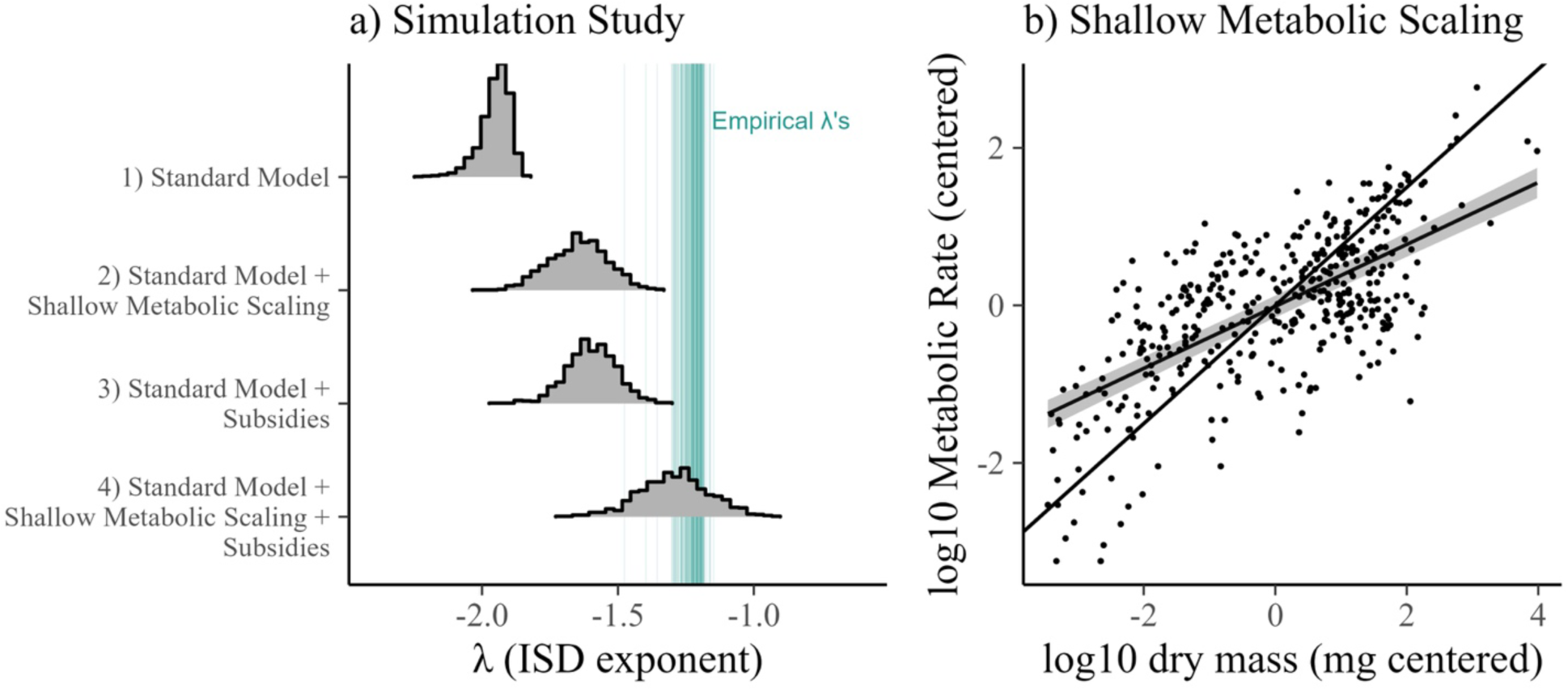
Empirical ISD’s in streams can only be explained by shallower metabolic scaling and subsidies. a) Simulations of λ from four conditions of the model: 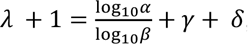, where *α* is the predator-prey mass ratio, *β* is trophic transfer efficiency, γ is the metabolic scaling exponent, and *δ* is the effect of ecological subsidies^32,34^. Histograms represent four simulations from this equation with different parameter inputs (Table S1), ranging from a Standard Model^20^ with ε fixed at 0.75 to model 4 that incorporates shallow metabolic scaling ε and ecological subsidies. Parameter inputs are described in Table S1. Only the model with subsidies and shallow metabolic scaling match the empirical lambdas. b) To test shallow metabolic scaling (*α*), we measured metabolic scaling in 414 macroinvertebrates from freshwater mesocosms. The regression line and shading show the median slope of 0.39 (95% CrI: 0.36 to 0.43), substantially shallower than the theoretical expectation of 0.75 (solid reference line), and in line with the value needed to match empirical lambdas in the simulation study.

An alternative solution is to modify Eq. 1 to include the influence of terrestrial subsidies to large organisms such as fish. Such subsidies (e.g., falling leaves and insects) are common in freshwater streams and help explain how streams can support more predators than expected from macroinvertebrate secondary production alone^33^. Terrestrial subsidies have been shown to cause a 0.2 to 0.5 unit increase in *λ* in freshwater streams^32,34^. Similarly, detrital subsidies from pelagic to benthic marine food webs led to shallower λ in benthic compared with pelagic food webs^19^. Adding subsidies to our simulations yielded λs that are still far away from the empirical median (simulated λ median of −1.6 instead of −1.2; Fig. 4a).

Only by relaxing the assumption of 0.75 metabolic scaling were we able to recapture the empirical distributions using simulations of Eq. 1 (Fig. 4a). The results suggest that λs of −1.2 can only be achieved by assuming γ ∼0.2 to 0.5, instead of 0.75, along with adding subsidy effects of 0.2 to 0.5. Surprisingly, we are not aware of any studies measuring γ for freshwater invertebrates or fish at community-level, though values of γ in the range of 0.2 to 0.5 have been reported for other marine invertebrates at species-level^35^. To test this further, we conducted a metabolic scaling experiment using 24 large outdoor freshwater mesocosms and a naturally colonizing community of macroinvertebrates. Strikingly, the experiment revealed γ to be ∼0.4 across a range of taxa, temperature, and predation scenarios (Fig. 4b), consistent with the simulation predictions explaining λ in North American freshwater streams (Fig. 4a).

Stream food webs deviate from the anticipated ISD λ value of ∼ −2, and exhibit metabolic scaling exponents that differ from the expected value of 0.75. Our findings endorse the existence of shared constraints that govern the size structure of geographically distributed stream food webs. These constraints likely stem from commonalities in the size scaling of metabolism^20^ and the trade-offs between the between the number of individuals at each size and the amount of energy flow they can sequester in stream ecosystems.

## Conclusions

The most important result of this work is the invariance of size spectra across a wide natural environmental gradient of temperature and resource supply in North American freshwater streams. The fact that this invariance occurs at macroecological scales and over years of repeated data collection suggest potential constraints in the ecological processes that generate size spectra in streams. The influence of temperature and resource availability on λ appears negligible, suggesting stable community size structure across extensive spatial scales, aligning with findings from previous studies^15^. Our results demonstrate a consistently higher proportion of large individuals across various environmental conditions.

Prior studies of the change in λ with temperature have predominantly focused on individual taxonomic groups (e.g., phytoplankton^25^, macroinvertebrates^27^, fish^36^) or restricted size ranges^36^. Furthermore, these investigations are confined spatially and temporally^4,25,37^, with distinct patterns emerging in different seasons (e.g., April vs. October), with many studies concentrating on relatively narrow intervals. Specifically, the most restricted range was approximately ∼4°C^25,36,37^ followed by ∼10 degrees^4,28^, and the widest being ∼15°C^27^.

We incorporate these various study characteristics into a comparison with the current results in Fig. 5. It shows the variation in reported change in λ with temperature, but also relatively small effect sizes as noted in macroinvertebrate communites^27^. These characteristics do not indicate shortcomings in the studies but underscore the challenges in extrapolating findings to comprehend the broader taxonomic, spatial, and temporal shifts in the size spectrum of ecological communities.

**Fig. 5.**
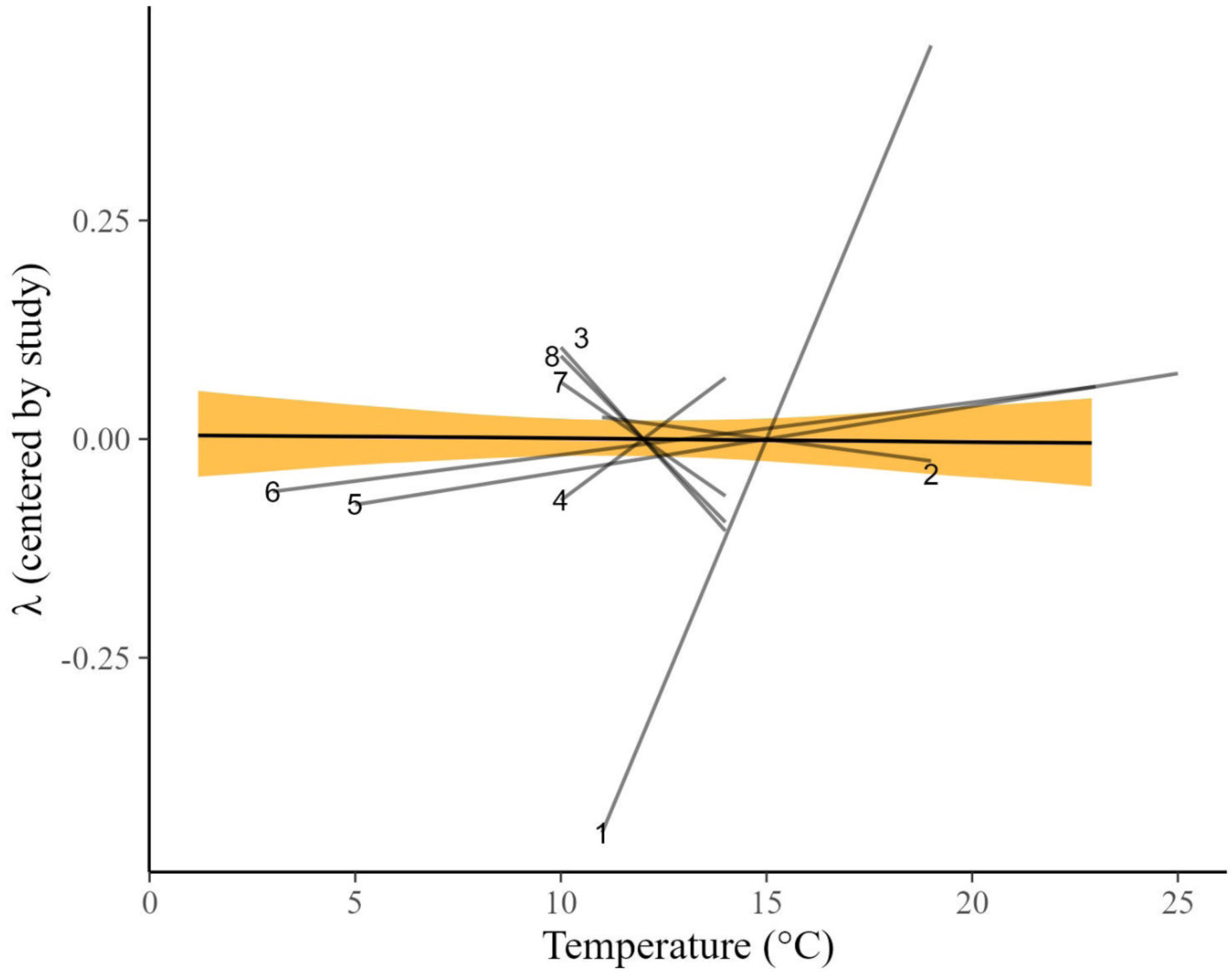
Variation in literature estimates of the ISD-temperature relationship compared to the current work. The current NEON-based analysis is in yellow. Individual lines show the change in lambda across 9 additional studies. Lambda on the y-axis is centered to the mean for each study to make the different approaches in reporting and estimating lambda comparable. The figure shows several large temperature ranges (∼15 degrees C), with most studies having relatively narrow ranges (∼5 C), with particular focus on temperate streams around 12 C. Lambda values were estimated figures, tables, or summary statistics in the corresponding study. 1^28^, 2^28^, 3^37^ (April estimate), 4^37^ (October estimate), 5^4^, 6^27^, 7^25^ (community), 8^25^ (phytoplankton only).

Indeed, when we re-ran our analyses on macroinvertebrates or fish separately, we found that λ declined with temperature for macroinvertebrates (Fig. S2b) and was more variable for fish (Fig. S2c), reconfirming previous studies on individual taxonomic groups^27,36^. An explanation for this is that the ISD for fish is not independent of invertebrates and vice versa, since they are trophically connected in size-structured food webs. Consequently, excluding either fish or invertebrates from the analysis results in the omission of the largest individuals (fish) and/or the most abundant small individuals (macroinvertebrates). This limitation may hinder the ability to scale taxonomically discrete size distributions up to the community level. Such influences are noticeable in marine food webs, where the presence or absence of macroinvertebrates has influenced conclusions drawn about the size distribution^38^.

Another possible explanation is that TTE, PPMR and metabolic scaling are also invariant across temperature and resources, since the ISD arises from these factors. This would be consistent with previous empirical research in marine food webs^39^, which observed the absence of systematic variations in PPMR and TTE across different temperature and resource conditions. Alternatively, as our simulations demonstrate (Fig. 4a), there are multiple combinations of TTE, PPMR, metabolic scaling, and subsidies that can generate the empirical λ values shown here. This implies that the variables driving λ can vary systematically with temperature without necessarily changing λ. For example, the biomass of larger consumers is expected to decline with increasing temperature^23^, which should reduce the PPMR. In addition, warmer temperatures can reduce TTE due to a reduction in the nutritional value of individual prey^31^. If warming has the same relative effect on log10(PPMR) and log10(TTE), then there would be no net change in their ratio, leading to constant λ. In contrast, unequal changes in the PPMR and TTE can yield steeper λ^40^ by limiting the amount of energy available to larger consumers.

Our study reveals that size spectra in freshwater streams remain consistent across temperature and resource gradients. Because our results occur in relatively unimpacted streams across broad temperature gradients, they likely reflect a macroecological pattern, but do not discount potential future effects of climate change^28^. In other words, comparing communities across spatial temperature gradients is different than observing an individual community respond to rapid environmental warming. The main challenge moving forward is understanding the fundamental principles behind this stability and forecasting future trends. Our approach not only aids in uncovering factors influencing larger-scale community processes but also holds promise for predicting ecosystem dynamics across different environments and scales.

## Methods

### Body Size Data

We analyzed size spectra using 2,468,683 individual body masses of fish and macroinvertebrates collected by the National Ecological Observatory Network (NEON) between 2016 and 2021. This included 64,940 measures from 157 fish species and 2,403,743 measures from 1,155 macroinvertebrates taxonomic groups (typically genus or species). The samples were collected once or twice per year at each of 23 sites, resulting in a total of 123 unique collection events.

### Macroinvertebrates

NEON collected macroinvertebrate data via fixed-area samplers (e.g., Surber/Core/Kicknet) and measured insect body lengths to the nearest mm along with estimates of their density (no/m^2^). The macroinvertebrate data are available as data product DP1.20120.001 (NEON 2023). While the samplers vary, all mesh sizes are the same (243 um). Measurements of macroinvertebrate lengths spanned from 1 to 86 mm, subsequently converted to dry mass employing published taxon-specific length-mass regressions^44^. These regressions included genus-specific species in some instances. After converting to dry mass, we excluded insects that were smaller than 0.0026 mg DM, as previous analyses showed these sizes to be under sampled^32^.

### Fish

Fish were collected from each site twice per year (typically) using 3-pass removal electrofishing. The fish data are available as data product DP1.20107.001 (NEON 2022). For each collection, the first 50 fish per taxon were measured for total length in mm and wet mass in mg^45^. The remaining fish were tallied as a bulk count per species (without mass measures). Using the three-pass depletion data, we estimated fish population density (no/m^2^) in each collection using a multinomial Poisson depletion model^46^. We specified the model in R using the *ubms* package^47^. The response variable was the number of fish caught per pass and the predictor variable was the collection id (*site*+*date*+*reach* unique ID). The model resulted in a population estimate for each collection, which we converted to no/m^2^ by dividing each estimate by the sampled area. We then multiplied that population estimate by the relative abundance of each fish species, resulting in an estimated density (no/m^2^) of each fish species in each collection. Finally, we merged those estimates with the dry mass measurements and resampled the dry mass measures with replacement, weighted by the relative abundance of each fish species (in units of number of individuals/10,000 m^2^). Weighting by no/10,000m^2^ instead of no/m^2^ was necessary to ensure that enough body sizes were sampled (i.e., no/m^2^ was typically <1). For each collection, we then summed the total number of individual body sizes, along with their density estimates, resulting in a dataset containing individual size estimates and their associated density for each collection.

### Combining fish and macroinvertebrates

Fish and macroinvertebrates were collected on different dates, with macroinvertebrates collected three times per year and fish collected twice. Therefore, to combine fish and macroinvertebrate samples, we limited the data to only collections that occurred within 30 days of each other. For example, if macroinvertebrates were collected on June 10 and fish collected on June 20, those samples were treated as one. If more than one sample was in this window (e.g., another fish collection on June 21), we included only the most recent collection. The resulting data set contains body sizes ranging nine orders of magnitude (0.003 to 200,000 mg) along with their densities. We used this dataset to estimate individual size distributions, total community biomass.

### Environmental Data

#### Temperature

To estimate mean annual stream temperature for each of the 22 sites, we obtained water temperature readings collected every four hours from 2016 to 2021 using the NEON data product DP1.20053.001 (NEON 2023b). We removed data that did not pass quality checks as noted by NEON. We also removed data that appeared unreasonably low (< −5°C) or high (> 50°C). Some data for Alaskan streams is missing when the water is frozen. For those data, we assumed a temperature of 0°C. The resulting dataset contained 93,930 temperature readings. To reduce the data size for modeling, we estimated the mean weekly temperature and modeled that as a function of date and site using a generalized additive model with a Gaussian likelihood and year as a varying intercept. Mean weekly temperature was centered prior to modeling. This approach allowed us to have a posterior distribution of temperature predictions on each day over three years. From that posterior distribution, we calculated the mean annual temperature and standard deviation (Table 1) for each site.

**Table 1.** Mean annual (± sd) values of temperature (°C) and gross primary production (GPP, gC/m^2^/y), along with standing stock organic matter (OM, AFDM/m^2^), at the 22 NEON sites in this study. Values are averaged over multiple years for temperature and GPP using sensor readings from NEON. For OM, values are calculated directly from benthic samples.

| Site | Temperature | GPP | OM |
| --- | --- | --- | --- |
| ARIK | $12 \pm 8$ | $1728 \pm 396$ | $5 \pm 4$ |
| BIGC | $10 \pm 5$ | $727 \pm 194$ | $4 \pm 2$ |
| CARI | $1 \pm 2$ | $353 \pm 44$ | $3 \pm 1$ |
| GUIL | $21 \pm 2$ | $4235 \pm 1573$ | $3 \pm 4$ |
| HOPB | $9 \pm 7$ | $541 \pm 198$ | $5 \pm 5$ |
| KING | $12 \pm 5$ | $1919 \pm 580$ | $92 \pm 126$ |
| LECO | $13 \pm 5$ | $1402 \pm 387$ | $3 \pm 1$ |
| MART | $8 \pm 4$ | $891 \pm 312$ | $5 \pm 1$ |
| OKSR | $1 \pm 3$ | $420 \pm 256$ | $4 \pm 2$ |
| REDB | $6 \pm 4$ | $821 \pm 326$ | $1 \pm 2$ |
| WALK | $14 \pm 3$ | $403 \pm 40$ | 46 |
| WLOU | $2 \pm 3$ | $351 \pm 88$ | $19 \pm 7$ |
| LEWI | $13 \pm 4$ | $4828 \pm 823$ | $377 \pm 418$ |
| TECR | $6 \pm 5$ | $350 \pm 182$ | $35 \pm 21$ |
| BLUE | $16 \pm 4$ | $7925 \pm 2536$ | $2 \pm 0$ |
| CUPE | $23 \pm 1$ | $3344 \pm 298$ | $1 \pm 1$ |
| MCDI | $13 \pm 7$ | $1573 \pm 1205$ | $5 \pm 8$ |
| MCRA | $7 \pm 4$ | $1719 \pm 308$ | $7 \pm 4$ |
| POSE | $12 \pm 7$ | $250 \pm 48$ | $12 \pm 13$ |
| PRIN | $17 \pm 7$ | $2479 \pm 557$ | $20 \pm 23$ |
| SYCA | $19 \pm 6$ | $11957 \pm 4169$ | $18 \pm 16$ |
| BLDE | $4 \pm 5$ | $907 \pm 270$ | $5 \pm 4$ |

#### Gross Primary Production

We estimated regimes of annual gross primary production (GPP, g C m^−2^ y^−1^) through a multistep ensemble model process (Supplemental Materials). Daily time series of NEON sensor-based data for temperature (DPI # 20053.001), stream discharge (DPI # DP4.00130.001), and oxygen concentration (DPI # DP1.20288.001) were checked for data quality flags and visually inspected to identify potential errant data. We then estimated daily GPP from cleaned time series with maximum likelihood estimation through 15 different methods using the ‘*streamMetabolizer*’ package^48^. Methods varied in respect to discharge– gas exchange, photosynthetic rate–light, and photosynthetic rate–temperature relationships. From this model suite, we first calculated root mean squared error (RMSE) based on pointwise differences between predicted and observed daily oxygen series. This model suite was reduced by removing models that yielded unreasonably high estimates based on mean daily GPP (>13,000 g C m^−2^ y^−1^), correlation coefficients >0.95 between reaeration and ecosystem respiration, and particularly poor fits based on RMSE. From this filtered model suite, we created an ensemble model weighted by relative RMSE. Finally, to develop a general model of the GPP regime at each site, we fit a hierarchical generalized additive model (GAMM) to ensemble estimated GPP by ‘day of year’ with ‘year’ treated as a random effect term. We summed the estimated ‘day of year’ fixed effect to quantify annual GPP for all sites. Annual GAMMs were fit with the ‘*brms*’ package^49^ with smoothing functions from the ‘*mgcv*’^50^.

#### Organic Matter

We estimated the standing stock of organic matter from 49 unsorted bulk benthic samples in the NEON biorepository (at least two samples per site)^51^. The samples were collected at the same time as macroinvertebrate samples, using the same collection techniques (i.e., Surber sampler, core sampler). We obtained the raw samples from NEON and first removed macroinvertebrates. We dried the remaining organic matter at 60 °C for >48 hours to bring to a constant mass before determining the initial mass. We then combusted samples at 500 °C for four hours before reweighing to determine total organic matter mass. Samples were scaled to areal mass (g m^−2^) based on sampler area.

### Metabolic scaling experiment

We conducted a mesocosm experiment, mirroring stream conditions, wherein factorial combinations of two temperature levels (natural vs. heated) and two predation regimes (with predation vs. without predation) were implemented. A sample of macroinvertebrates was collected from each tank 30 days after water first flowed through them. This timeframe was chosen as it allows for ongoing changes in community composition, with the size spectra expected to stabilize around an equilibrium within the specified experimental period of 30 days.

To determine the general metabolic scaling of the community, *M*^α^, we computed the metabolic rate based on individual macroinvertebrates gathered from each tank. Using oxygen consumption as an indicator for standard metabolic rate, we followed established methods^52^. Macroinvertebrates were subjected to a 24-hour starvation period prior to measuring oxygen consumption to eliminate the energetic costs of specific dynamic action and minimize the accumulation of excretory products.

Subsequently, each individual was placed in a transparent glass vial containing 20 mL of oxygenated, filtered water. After one hour, we measured the reduction in oxygen concentration within each vial using oxygen sensors (e.g., PSt3, PreSens, Regensburg, Germany) and a portable oxygen meter equipped with a fiber-optic cable (e.g., Fibox 4 trace, PreSens, Regensburg, Germany). Corrections were made for changes in oxygen concentrations in control vials (without macroinvertebrates), and individual metabolic rates were standardized to milligrams of oxygen consumed per liter per hour (mg L^−1^ hour^−1^).

Furthermore, after respirometry measurement, each individual was dried and measured for individual mass on a microbalance, enabling the assessment of the relationship between metabolic rate and body size.

### Data Analysis

To examine how size spectra varied as a function of temperature and resources, we used a Bayesian generalized linear mixed model with a truncated Pareto likelihood. A description and justification of this modelling approach for ISD’s is given in^53^. The model structure was:

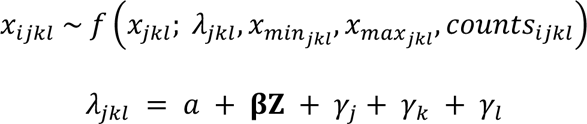

where *x_ijkl_* is the *i*^th^ body size from sample *j* in site *k* and year *l*. The likelihood *f(…)* is a truncated Pareto with a single free parameter *λ_jkl_*, the exponent of the ISD. *x_minjkl_* and *x_maxjkl_* are the minimum and maximum body sizes in each sample, site, and year. Each body size has a corresponding density in units of number per m^2^, represented by counts*_ijkl_*, as described in^54^. *λ_jkl_* is modeled as a linear function of an intercept *α* and **βZ** represents the single, two, and three-way interactions of the **Z** predictors of mean annual temperature, mean annual GPP, and standing stock organic matter. All predictors were standardized as z-scores prior to fitting. Varying intercepts are included for individual sample (*γ_j_*), site *γ_k_*), and year (*γ_l_*). To improve sampling efficiency, the varying intercepts were modeled using non-centered parameterization, which is excluded here for clarity, but is present in the Stan model code: https://github.com/jswesner/neon_size_spectra-slim.

Priors for the intercept were Normal(−1.5, 0.2), chosen based on a previous analysis of ISD values in NEON streams^27^. Priors for each *β* parameter were set to Normal(0, 0.1). Priors for the *q* varying intercepts were Normal(0, α_q_), with each α_q_ hyperprior set to Exponential(7). These priors were chosen based on prior predictive simulation^55^ to center the prior probabilities of λ between about −2 to −1 while still allowing probabilities at very large (e.g., 1) or small values (e.g., −4) (Fig. S3).

To test the metabolic scaling relationship, we used a gaussian linear mixed model with log_10_ respiration (centered at zero) rate as the response variable, log_10_ dry mass (centered at zero), heat (presence/absence), and fish (presence/absence), and their interactions as predictor variables, and mesocosm tank as a varying intercept. Priors for the intercept were Normal(0, 1). The prior for the marginal slope between log_10_ respiration and log_10_ dry mass was Normal(0.75, 2), and all other priors for the regression parameters were Normal(0, 1).

We fit all models in *rstan* via the *brms* and *isdbayes* packages in R. Each model had 4 chains with 2000 iterations, where the first 1000 were discarded as warm-up. Model fit was checked using posterior predictive checks and prior influence was checked using prior predictive simulation.

## Acknowledgments

This material is based upon work supported by the National Science Foundation under Grant Nos. 2106067 to JSW and 2106068 to JRJ. The National Ecological Observatory Network Biorepository at Arizona State University provided samples for organic matter collected as part of the NEON Program. The use of vertebrates was approved under the University of South Dakota Institutional Animal Care and Use Committee (#03-03-18-21). Computations supporting this project were performed on High Performance Computing systems at the University of South Dakota, funded by NSF Award OAC-1626516.

## Supplementary Information

**Table S1.** Parameters of four simulation scenarios of factors that explain lambda: Predator Prey Mass Ratio (PPMR), Trophic Efficiency (TE), Metabolic Scaling Coefficient (MS), and Ecological Subsidies. The mean, median, and sd are derived from 10,000 simulations for each parameter with the following probability distribution functions in R: PPMR = rlnorm(10000, 12, 3), TE = rbeta(10000, 7, 60), MS = rbeta(10000, 12, 25), Subsidies = rbeta(10000, 14, 15). In models 1-3, at least one parameter is fixed (e.g., MS is assumed to be 0.75 in models 1 and 3).

| Parameter | Model | Constant | Mean | Media<br>n | SD |
| --- | --- | --- | --- | --- | --- |
| PPMR ($\alpha$) | 1) Standard Model | No | $10^7$ | $10^5$ | 4E+08 |
| TE ($\beta$) | | No | 0.1 | 0.1 | 0.04 |
| MS ($\varepsilon$) | | Yes | 0.75 | 0.75 | 0 |
| Subsidies ($\delta$) | | Yes | 0 | 0 | 0 |
| PPMR ($\alpha$) | 2) Standard Model + Shallow<br>Metabolic Scaling | No | $10^7$ | $10^5$ | 4E+08 |
| TE ($\beta$) | | No | 0.1 | 0.1 | 0.04 |
| MS ($\gamma$) | | No | 0.4 | 0.4 | 0.09 |
| Subsidies ($\delta$) | | Yes | 0 | 0 | 0 |
| PPMR ($\alpha$) | 3) Standard Model + Subsidies | No | $10^7$ | $10^5$ | 4E+08 |
| TE ($\beta$) | | No | 0.1 | 0.1 | 0.04 |
| MS ($\gamma$) | | Yes | 0.75 | 0.75 | 0 |
| Subsidies ($\delta$) | | No | 0.4 | 0.4 | 0.08 |
| PPMR ($\alpha$) | 4) Standard Model + Shallow<br>Metabolic Scaling + Subsidies | No | $10^7$ | $10^5$ | 4E+08 |
| TE ($\beta$) | | No | 0.1 | 0.1 | 0.04 |
| MS ($\gamma$) | | No | 0.4 | 0.4 | 0.09 |
| Subsidies ($\delta$) | | No | 0.4 | 0.4 | 0.08 |

**Fig. S1.**
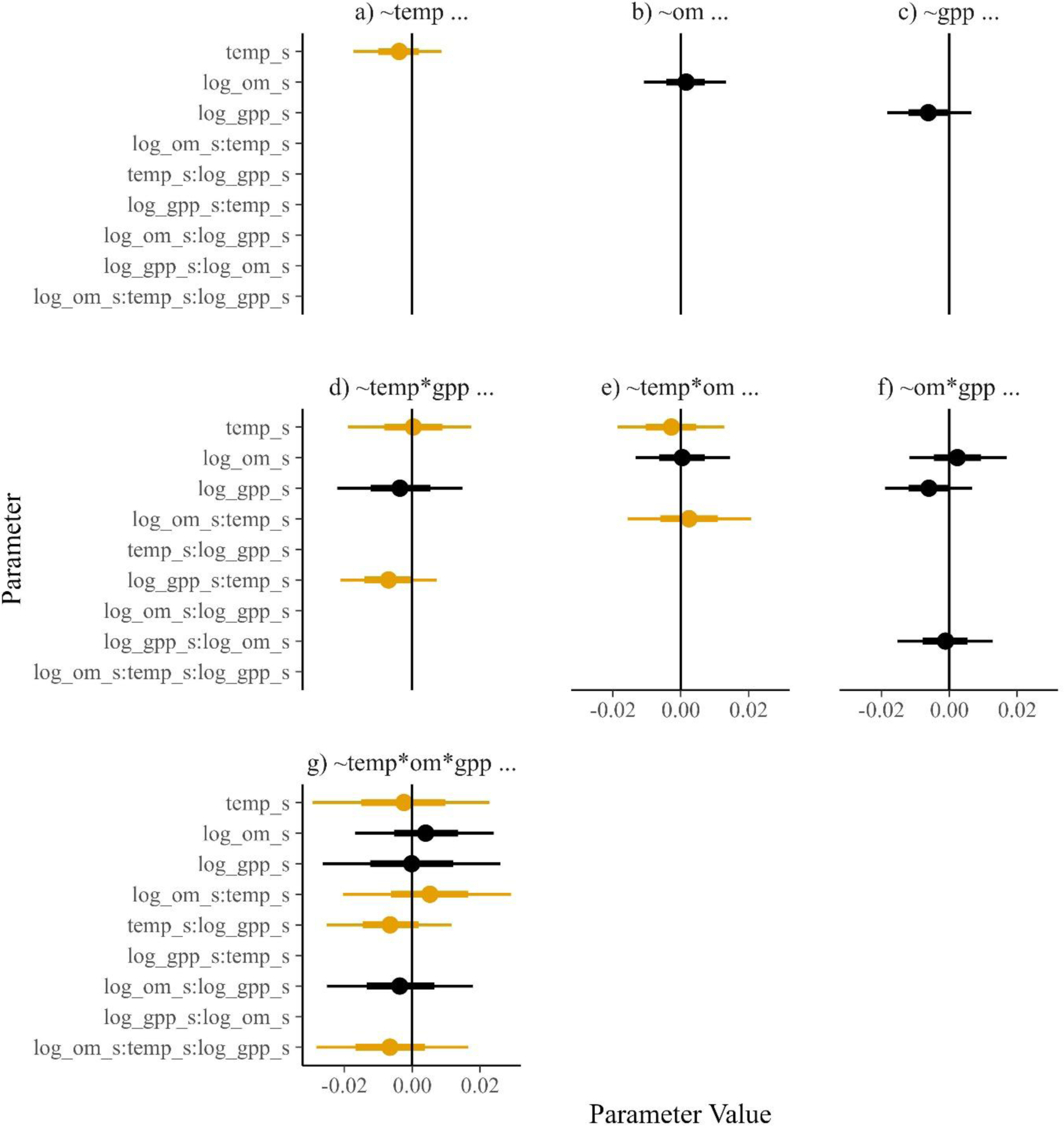
Posterior distributions of fixed effects parameters from seven models estimating change in λ across environmental predictors. Models contain either univariate (a-c), two-way interactions (d-f), or a three-way interaction (g) of temperature (“temp”), gross primary production (‘gpp”) or organic matter standing stock (“om”). All variables were standardized with z-scores prior to analysis. GPP add organic matter were log10 transformed prior to standardization. Parameters containing temperature are highlighted in yellow. All models contained variating intercepts of year, site, and sample, which is abbreviated with elipses (…). The median, 50, and 95% Credible Intervals are shown by the dot, thick bar, and thin bar, respectively.

**Figure S2.**
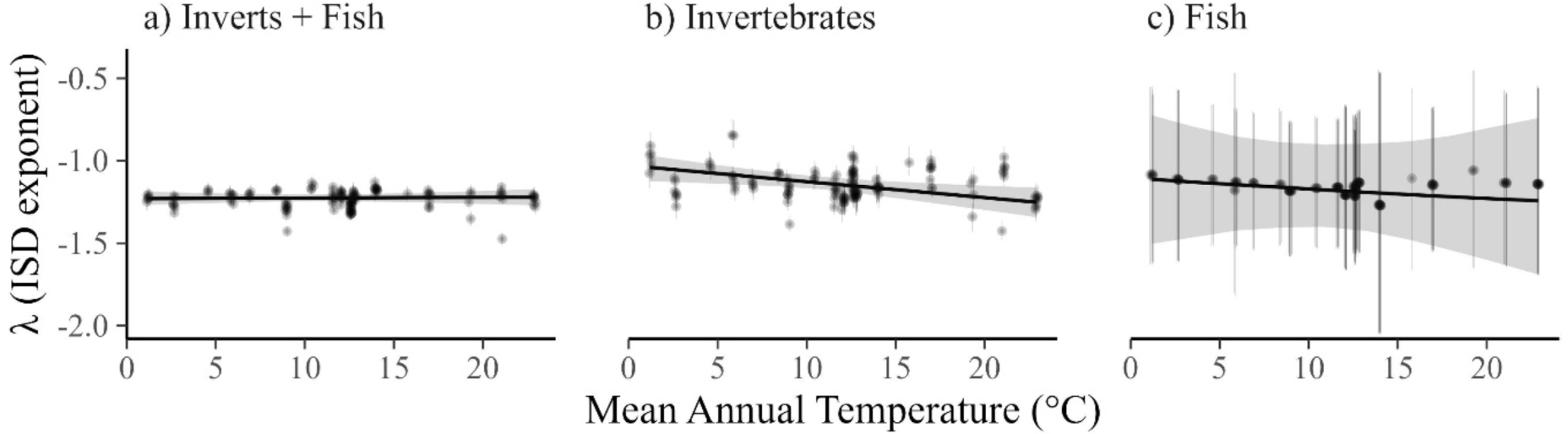
Community-wide results differ from individual taxa results. a-c) Invariance of the ISD for the community (a) does not hold when only analyzed for invertebrates (b). For fish (c), the level of replication (number of individual fish per sample) is lower, and the relationship more uncertain than a or b. Lines are posterior medians and shading is the 95% Credible interval. Dots in a-c are sample-specific lambdas predicted from varying intercepts in each model.

**Figure S3.**
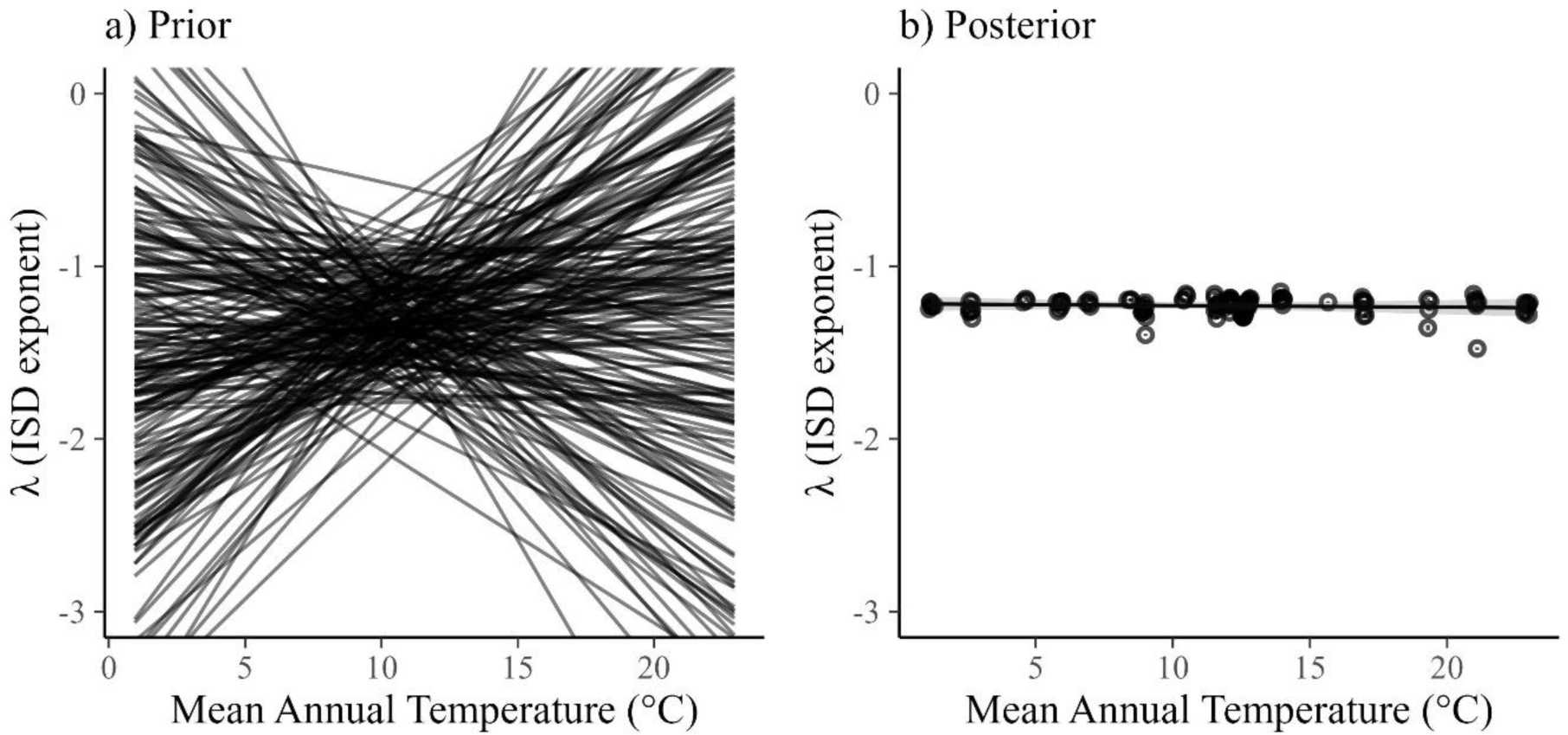
Two-hundred simulations from the prior predictive distribution. (a) compared to the fitted posteriors (b). Both models show the relationship between λ and temperature with GPP and organic matter set to their median values. We checked the implications of our priors using prior predictive simulation. The result is shown in Fig. S3, indicating that the priors largely limit λ to values between ∼-2 to −1, but allow for a wide range of possible relationships with mean annual temperature. By comparison, the posterior (Figure S1b) remains in a much more constricted space. The difference between the prior and posterior is an index of how much information was learned from the data.

